# Limits and potential of combined folding and docking using PconsDock

**DOI:** 10.1101/2021.06.04.446442

**Authors:** Gabriele Pozzati, Wensi Zhu, Claudio Bassot, John Lamb, Petras Kundrotas, Arne Elofsson

## Abstract

In the last decade, de novo protein structure prediction accuracy for individual proteins has improved significantly by utilising deep learning (DL) methods for harvesting the co-evolution information from large multiple sequence alignments (MSA). In CASP14, the best groups predicted the structure of most proteins with impressive accuracy. The same approach can, in principle, also be used to extract information about evolutionary-based contacts across protein-protein interfaces. However, most of the earlier studies have not used the latest DL methods for inter-chain contact distance prediction. This paper introduces a fold-and-dock method, PconsDock, based on predicted residue-residue distances with trRosetta. PconsDock can simultaneously predict the tertiary and quaternary structure of a protein pair, even when the structures of the monomers are not known. The straightforward application of this method to a standard dataset for protein-protein docking yielded limited success. However, using alternative methods for MSA generating allowed us to dock accurately significantly more proteins. We also introduced a novel scoring function, PconsDock, that accurately separates 98% of correctly and incorrectly folded and docked proteins. The average performance of the method is comparable to the use of traditional, template-based or *ab initio* shape-complementarity-only docking methods. However, no a priori structural information for the individual proteins is needed. Moreover, the results of conventional and fold-and-dock approaches are complementary, and thus a combined docking pipeline could increase overall docking success significantly. PconsDocck contributed to the best model for one of the CASP14 oligomeric targets, H1065.

## Introduction

Protein structure is crucial for our understanding of biological function. However, experimentally determining the structure of a protein is still time-consuming and expensive. Therefore, computational methods will be the only method to determine the structure of most proteins in the foreseeable future. Until recently, the only method to reliably predict the structure of a protein was to model it using a homologous template. However, reliable templates are not available for close to half the residues in the human proteome [1].

For several decades the prediction of protein structure directly from sequence information has been an unachievable dream. However, that changed about a decade ago when improved methods using co-evolution achieved sufficient residue contact information to predict the structure of many proteins [2,3]. Later, deep learning [4,5] and prediction of residue-residue distances provided further improvements [6,7]. Today this means that for many, if not most, individual proteins, it is possible to accurately predict the structure of its folded domains [8]. Recently, Deepmind demonstrated at CASP14 that using an end-to-end learnable approach, high-quality prediction of almost all protein domains is already feasible today (although not generally available).

In principle, the same type of methods used for predicting the structure of a single protein can predict the interaction between two proteins [9,10]. However, there is one fundamental difference: it is necessary to create paired alignments to identify the interaction between two proteins, i.e. identifying what pairs of proteins interact in the same manner. The identification of interacting pairs is assumed to be relatively easy for pairs of proteins that both only contain a single homolog in a set of genomes, but when multiple paralogs exist -the exact pairing is difficult [11].

Proteins do, however, not act alone. They function by interacting with other proteins and other molecules. Protein interaction can vary in nature from stable interaction present in small and large protein complexes to transient interactions often used for regulation. Experimentally the study of stable protein interactions can be done using various techniques. Structural determination methods, including crystallography and Cryo-EM electron microscopy, can solve the structure of protein complexes, while other methods can be used to identify that two proteins interact without obtaining detailed structural information.

Prediction of protein interactions has been an even more significant challenge than predicting the structure of individual proteins. Many different techniques have been developed, but in short, they can be divided into four categories: (i) docking primarily based on shape complementarity [12], (ii) template-based modelling [13], and (iii) flexible docking [14,15]. Various energy functions have also been used to improve the identification of correct docking poses [16]. In addition, co-evolution-based methods have also been used to predict the structure of complexes [9,17].

Benchmarks have been developed to elucidate the advantages and disadvantages of different docking methods [18]. Shape complementarity works excellently on native complexes, but the accuracy drops fast when using the structures of unbound complexes and even further if models of the proteins are used [19,20]. Template-based modelling works excellently if a complex with significant sequence identity exists in PDB but does not work for novel complexes[21,22].

Successful DCA based methods to predict protein-protein interactions preceded the large-scale prediction of single proteins by predicting the bacterial two-component signalling in 2009 [17]. These methods were then extended to a handful of other complexes by several groups [9,10]. However, it is still unclear how generally applicable these methods are, but the potential to vastly increase the space of known protein-protein interactions should lie in using some type of co-evolution based methods. The computational cost limits flexible docking, but a fold-and-dock protocol [23] based on coevolution does not require an exact structure of the two individual proteins.

In addition to determining the structure of a protein complex, it is also crucial to determine which proteins interact. However, protein-protein interaction is not an easily defined entity. It might include anything from proteins regulating the expression of genes to proteins strongly bound to each other in a large molecular machine. Several interaction databases exist [24,25], and methods, including co-evolution based methods [26], to predict interactions have been developed.

Here, we examine if it is possible to simultaneously fold and dock [23] two proteins by using coevolutionary information and not only dock them. In addition, we use one of the best methods (trRosetta) instead of DCA[2] based methods to predict intra- and inter-chain distances. One advantage of a fold-and-dock methodology is that it is not dependent on the availability of individual structures and should therefore be less sensitive to structural rearrangements upon binding. The disadvantage is that obviously, there are many more degrees of freedom in the system. We find that for several cases, it is possible to fold and dock the dimer simultaneously accurately. Although the success rate is low (<10%), this is comparable to the accuracy of other docking methods, which utilises the structure of both individual proteins. In addition, the methods are complementary.

## Results

The protocol used here starts from two multiple sequence alignments, created by searching with jackhmmer [27] against all complete proteomes from UniProt [28]. After that, a combined multiple sequence alignment is created by including the top paired hit from each proteome. It should be noted that the depth of the combined multiple sequence alignment is often significantly smaller than for the individual proteins. In addition, a few alternative methods both for generating the alignments and selecting the sequences were tried. These are discussed below. Next, twenty Glycine residues were added to separate the two sequences in the combined multiple sequence alignment. The combined alignment can be created in two different orientations, A-B vs B-A, and we have tried to use both combinations.

Next, the combined multiple sequence alignment has been adopted to predict distances and angles with trRosetta [29]. These are then used as input to Rosetta or CNS [30] to fold and dock the two proteins.

Below, we will discuss when this methodology works, when it fails, compare the performance of different alignments, and compare the performance with other docking techniques, and finally introduce a score, PconsDock, which accurately can be used to distinguish successful and unsuccessful docking attempts.

### Example of successful fold and dock

First, we demonstrate that the algorithm can accurately fold and dock a pair of proteins in at least one case. Figure 1 presents one successful example of the fold-and-dock protocol for the human protein complex between NOT1 MIF4G and CAF1 (PDB: 4gmj)[31]. The prediction is built on an alignment containing 1189 sequences (Meff=523) created by three iterations of jackhmmer[27] and an E-value cutoff of 10^−3^ against all reference proteomes in UniProt[28]. Visually, it can be seen that the intra-chain distance maps are similar and most intra-chains contacts are predicted accurately (PPV>0.90 for both chains), resulting in well-folded models of both chains (TM-score >0.8 for both). In total, 139 out of 287 inter-chain contacts are accurately predicted (287 contacts predicted with a PPV of 49%). The final docked model is also accurate (dockQ score 0.42). However, as we will show below, unfortunately, many models are not as easy to model as 4gmj. To test the performance of the algorithm, we have, therefore, used 222 heterodimeric protein pairs from dockground 4.3 [18,28].

**Figure 1:**
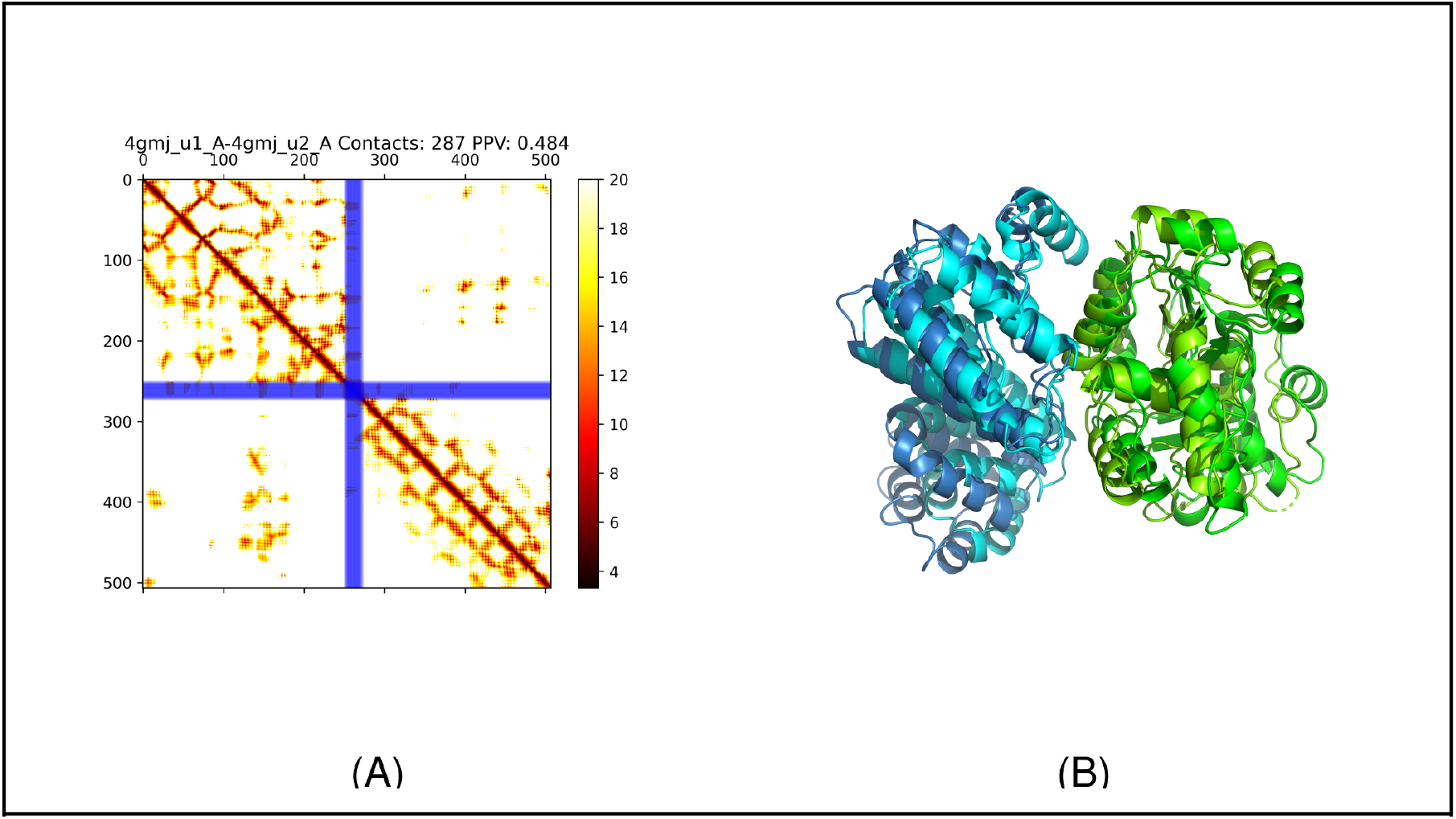
A) Predicted (lower triangle) and actual (upper triangle) distance map of the protein 4gmj. The two blue stripes represent the poly-G linker between the two chains. The title shows that 287 interchain contacts are predicted and that 48.4% of these are correct. B) Real (dark colours) and modelled (light colours) structure of the protein 1vrs. The accuracy of the models is good, dockQ score 0.42, and the TM-scores for the two chains are 0.82 and 0.85, respectively.

**Figure 2:**
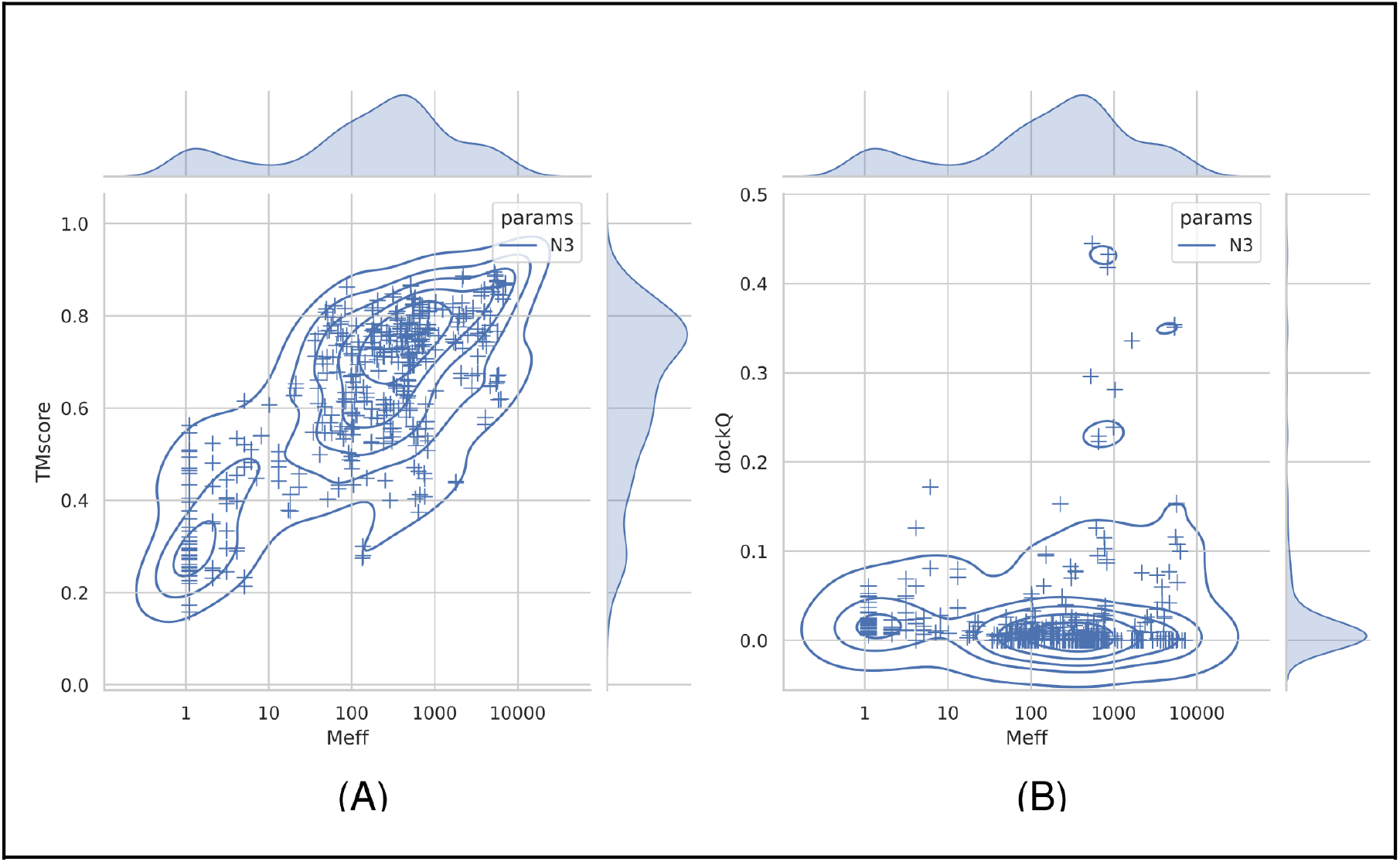
Performance of the fold-and-dock methodology versus the size of the joint alignments. Average TM-score of the two chains (A) and dockQ scores (B) plotted against the size of the multiple sequence alignment used to predict the contacts.

### Modelling accuracy depends on the size of the MSA; docking performance does not

The Dockground heterodimeric dataset was used to test the performance of the fold-and-dock methodology. First, we examined the dependence of the size of the multiple sequence alignment on the performance. It can be seen that the average TM-score for both chains is increasing with the size of the combined alignment, Figure 1. At a depth of 100 sequences, the average TM-score is over 0.6, indicating that about 100 effective sequences are in most cases sufficient to obtain the fold of a protein.

Next, we examined the quality of the predicted dimers, Figure 1B. A few models are docked correctly (dockQ score >0.23). However, most protein pairs are not accurately docked (dockQ score close to 0). The average dockQ score is only 0.02. Further, there is no apparent increase in the docking quality with more sequences in the MSA. What distinguishes the handful of docked models accurately is that they all seem to have between 100 and 1000 effective sequences in the merged alignments. However, it is also clear that many other protein pairs have MSAs of the same size but are not correctly docked. Next, we will examine how the docking is affected by the use of alternative alignments.

### Different alignments sometimes produce better models

We examined different cutoffs, different minimum coverage of the alignments to be included, and a different number of predictions to be included. We also tried to use a reciprocal best hits approach, i.e. only include proteins if they are orthologs, as described elsewhere [32]. In Figure 3, the folding and docking results for a selected subset of approaches can be seen, and for additional ones in Table 1. It can be noted that we also tried several other combinations, including the merging of predictions from alternative alignments, but none of these provided significant improvements, and for simplicity, we, therefore, focus on the methods used in Table 1.

**Table 1:** Overview of alignment methods used in this study (top rows) as input to the fold and dock protocol or alternative docking methods and their performance (Number of correct docked proteins, average dockQ score, and average TM-score.

| Name | Options/comments | No Acc. | <dockQ> | <TM> |
| --- | --- | --- | --- | --- |
| Best | Best (highest dockQ) prediction of methods marked with * | 15 | 0.069 | 0.57 |
| PconsDock | The top-ranked prediction of methods marked with * ranked by PconsDock | 10 | 0.040 | 0.58 |
| N1* | One iteration against reference proteomes with E=10<sup>-10</sup> cutoff | 6 | 0.027 | 0.55 |
| N3* | Three iterations against reference proteomes with E=10<sup>-3</sup> cutoff | 5 | 0.022 | 0.63 |
| N5 | Five iterations against reference proteomes with E=10<sup>-3</sup> cutoff | 3 | 0.019 | 0.63 |
| N1-minprob25* | Same as N1 but including all contacts with probability over 0.25 | 7 | 0.038 | 0.54 |
| N3-minprob25* | Same as N3 but including all contacts with probability over 0.25 | 5 | 0.032 | 0.63 |
| N1-cov50 | Same as N1 but only using sequence covering 50% of the joint alignment | 5 | 0.028 | 0.53 |
| N3-cov50 | Same as N3 but only using sequence covering 50% of the joint alignment | 6 | 0.024 | 0.58 |
| N1-cov90* | Same as N1 but only using sequence covering 90% of the joint alignment | 5 | 0.028 | 0.53 |
| N3-cov90* | Same as N3 but only using sequence covering 90% of the joint alignment | 6 | 0.024 | 0.58 |
| rbh-JH | Same as N3 but only reciprocal best hits using Jackhmmer | 3 | 0.018 | 0.56 |
| N3-merged | Same as N3, but distances for each chain predicted using complete alignments of each chain to predict intra-chain contacts. | 4 | 0.025 | 0.72 |
| N3-pdb | Same as N3 but contacts for each chain from the PDB file. | 5 | 0.019 | 0.92 |
| N3-bacteria | One iteration against bacterial reference proteomes with E=10<sup>-3</sup> cutoff | 3 | 0.022 | 0.47 |
| N1-all | One iteration against **all** proteomes with E=10<sup>-3</sup> cutoff | 5 | 0.023 | 0.60 |
| N3-all | Three iterations against **all** proteomes with E=10<sup>-3</sup> cutoff | 5 | 0.022 | 0.63 |
| RaptorX | Inter-chain contacts predicted by the RaptorX web server, intra-chain from PDB files. | 3 | 0.031 | 0.53 |
| Alternative methods |  |  |  |  |
| gramm | Shape-complementarity docking scored by the AACE18 potential | 16 | 0.081 | 1.0 |
| gramm-contact | Shape-complementarity docking scored by predicted contacts by trRosetta | 8 | 0.045 | 1.0 |
| gramm-raptorX | Shape-complementarity docking scored by contacts predicted by RaptorX | 6 | 0.051 | 1.0 |
| TMdock | Template-based docking, ignoring all templates with E-values < 10<sup>-5</sup> | 4 | 0.024 | 1.0 |

**Figure 3:**
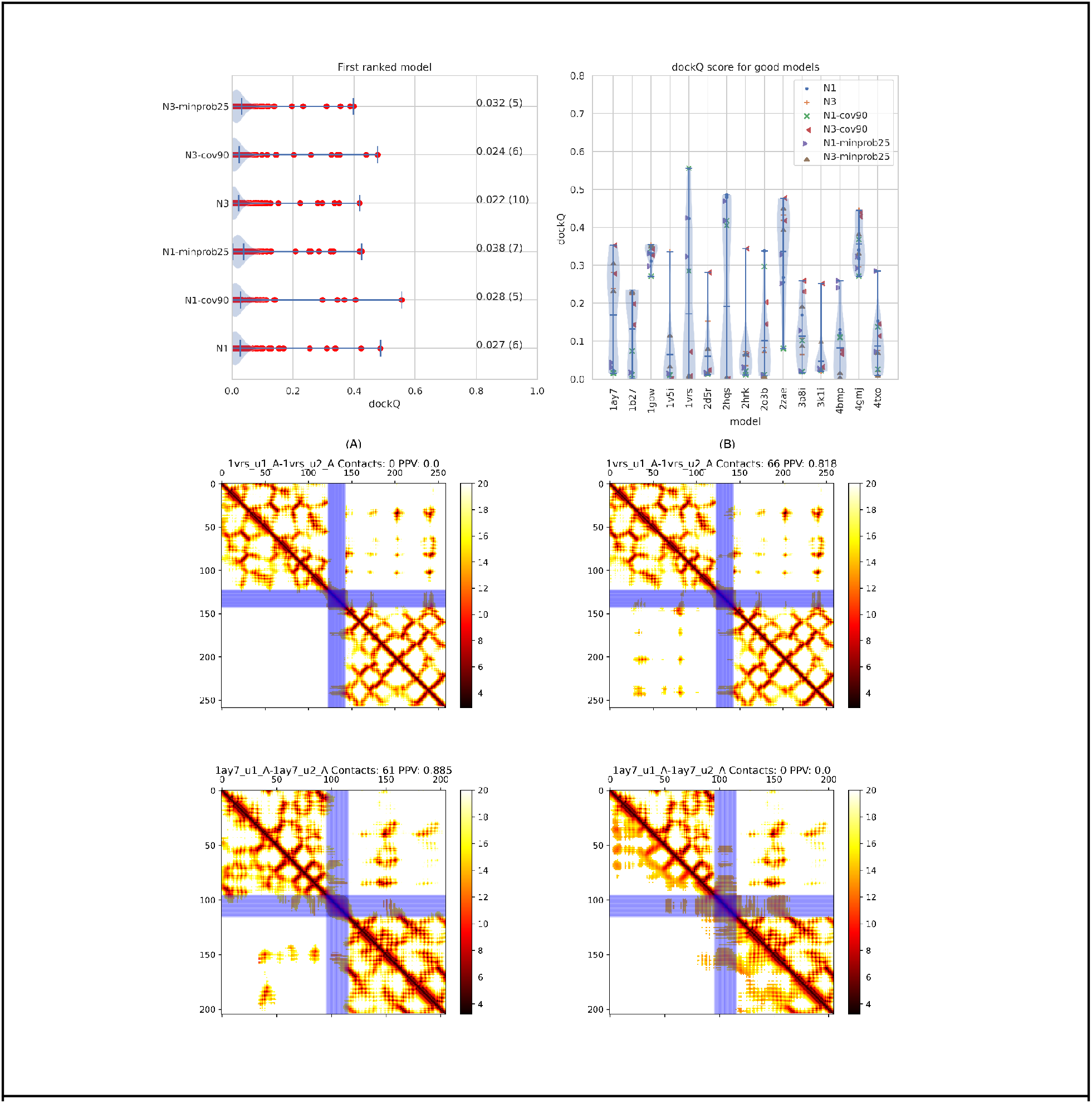
Results of PconsDock using different alignments. A) dockQ scores for all models using six different alignments (see Table 1). B) The dockQ scores for the 15 proteins where at least one of these six alignments produce an acceptable model (dockQ>0.23). C-F) Predicted contact maps for two proteins (1ay7 and 1vrs) plotted as described in Figure 1, using either (C, E) one iteration or (D, F) three iterations of Jackhmmer.

First, it can be noted that in some cases, one alignment methodology provides better contact maps than another. In Figure 3C-D, the contact maps of N- and C terminus of redox catalyst DsbD (PDB:1vrs) [33] using one (Figure 3C) or three (Figure 3D) iterations of jackhmmer searches are shown. When using three iterations, 66 contacts are predicted, and 82% of these are correct. In contrast, when using one iteration, zero inter-chain contacts are predicted. The opposite can be noted for the RNase Sa complex with Barstar (PDB:1ay7)[34], where one iteration makes a much better distance map than using three iterations, Figure 3E-F.

In Figure 3A, it can be seen that six different alignments methods roughly produce the same number of correctly docked models (three to seven). A similar trend can be seen for more methods in Table 1. Further, it is not always the same method that produces the best model, see Figure 3B. Here, it can also be seen that for most of the 15 models where at least one method produces a good model, there are only a few of the methods that produce a good model. The exceptions are 1gpw, 2zae, and 4gmj, where most methods produce good docking results, indicating that the fold-and-dock methodology could be improved if there was a methodology to identify the best way to generate the multiple sequence alignment.

### Comparison of Docking Protocols

We have also developed a CNS based method to fold-and-dock two proteins, named pyconsFold [35]. The advantage of this method is that it is about ten times faster than trRosetta, and the docking results are similar, Figure 4A. However, the quality of the independent proteins is less accurate, average TM-score 0.44 vs 0.63, Figure 4B. We also used RaptorX to predict inter-chain contacts using the webserver [36]. These results are also in line with the other results, and the docking results are not better than those obtained by trRosetta.

**Figure 4:**
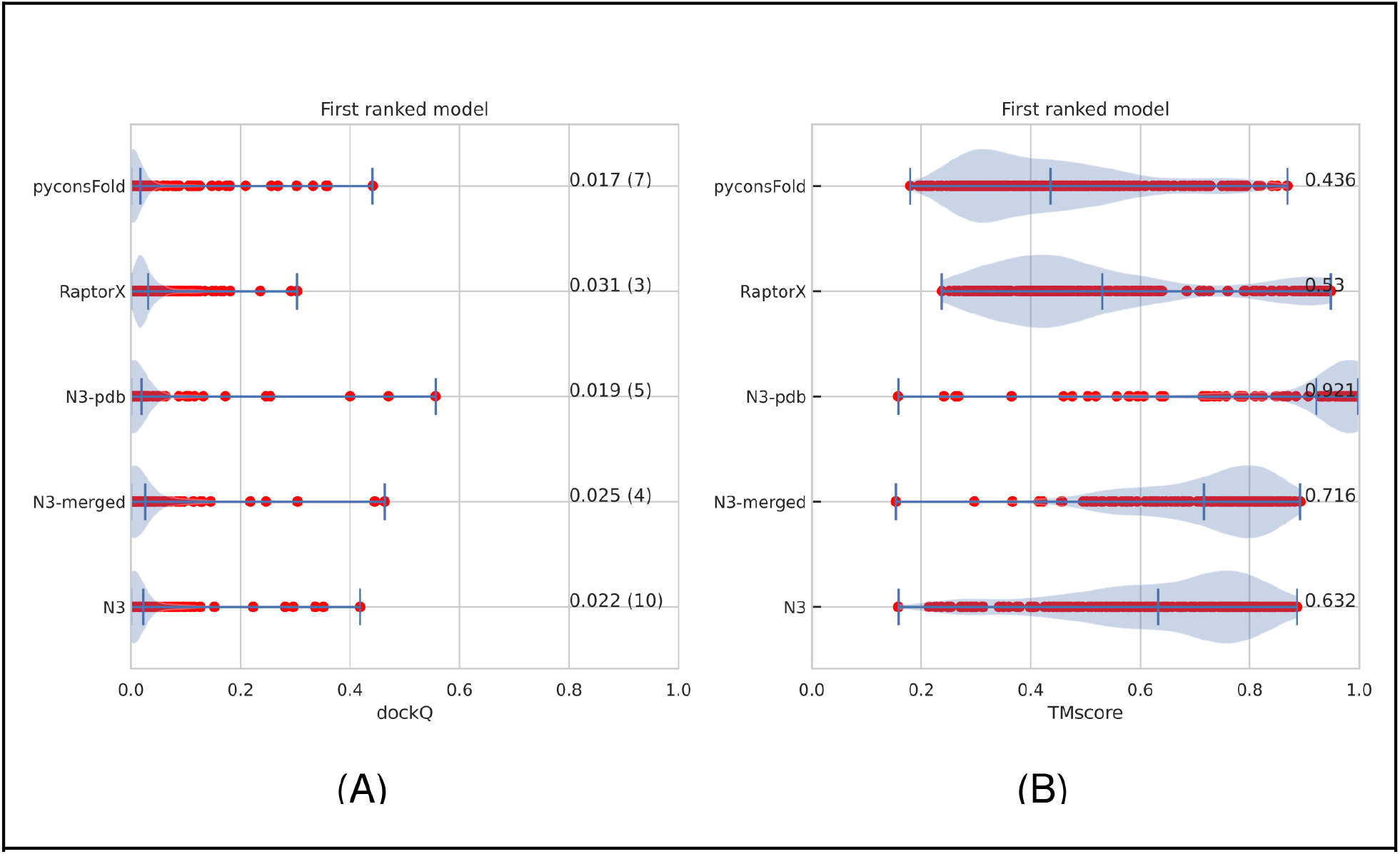
Results of alternative docking and folding protocols, average TM-score of the two chains (A) and dockQ scores (B). The default (N3) performance is compared withpyconsFold (uses the pyconsFold program instead of Rosetta), RaptorX (uses inter-chain contacts predicted by RaptorX instead of distances from trRosetta), RaptorX and N3-pdb use the intra-chain distances from the native structures, and N3-merged uses intra-chain distances predicted by the full alignments for each chain independently.

So far, we have used the same merges multiple sequence alignment to fold and dock the protein pairs. However, this is not the optimal way to use the information in the multiple sequence alignment for folding the individual chains. Instead, one can use the complete multiple sequence alignment for each of the two chains to predict the two intra-chain distance maps and then use these. Figure 4 shows that including more accurate intra-chain constraints improves the modelling of the individual chains, the average TM-score increases from 0.63 to 0.72, but the docking does not improve. Alternatively, it is also possible to use the distance from the structures, if available, of the individual proteins for the folding. Using this information improves the TM-score to 0.92, but the average docking results are not improved.

## Discussion

So far, we have shown that for 15 out of the 222 heterodimeric models in dockground 4.3. It is possible to create an acceptable (dockQ>0.23) model using predicted distances and a fold-and-dock protocol. However, no single alignment method does more than seven, i.e. if we can identify the optimal alignments for each target, it would be possible to improve the performance. Therefore, we first set out to identify factors that separate correct and incorrect models.

### Features separating correct and incorrect models

Most protein pairs can not be docked correctly, and for the few that do work, only a subset of the alignments work. What are the significant factors that distinguish the successful and unsuccessful cases? Figure 5 shows some features with some capacity for separation correct (dockQ>0.23) and incorrect models by plotting density plots. Note that there are many more incorrect models, so for comparison, we analyse the frequency of the features and not the numbers.

**Figure 5:**
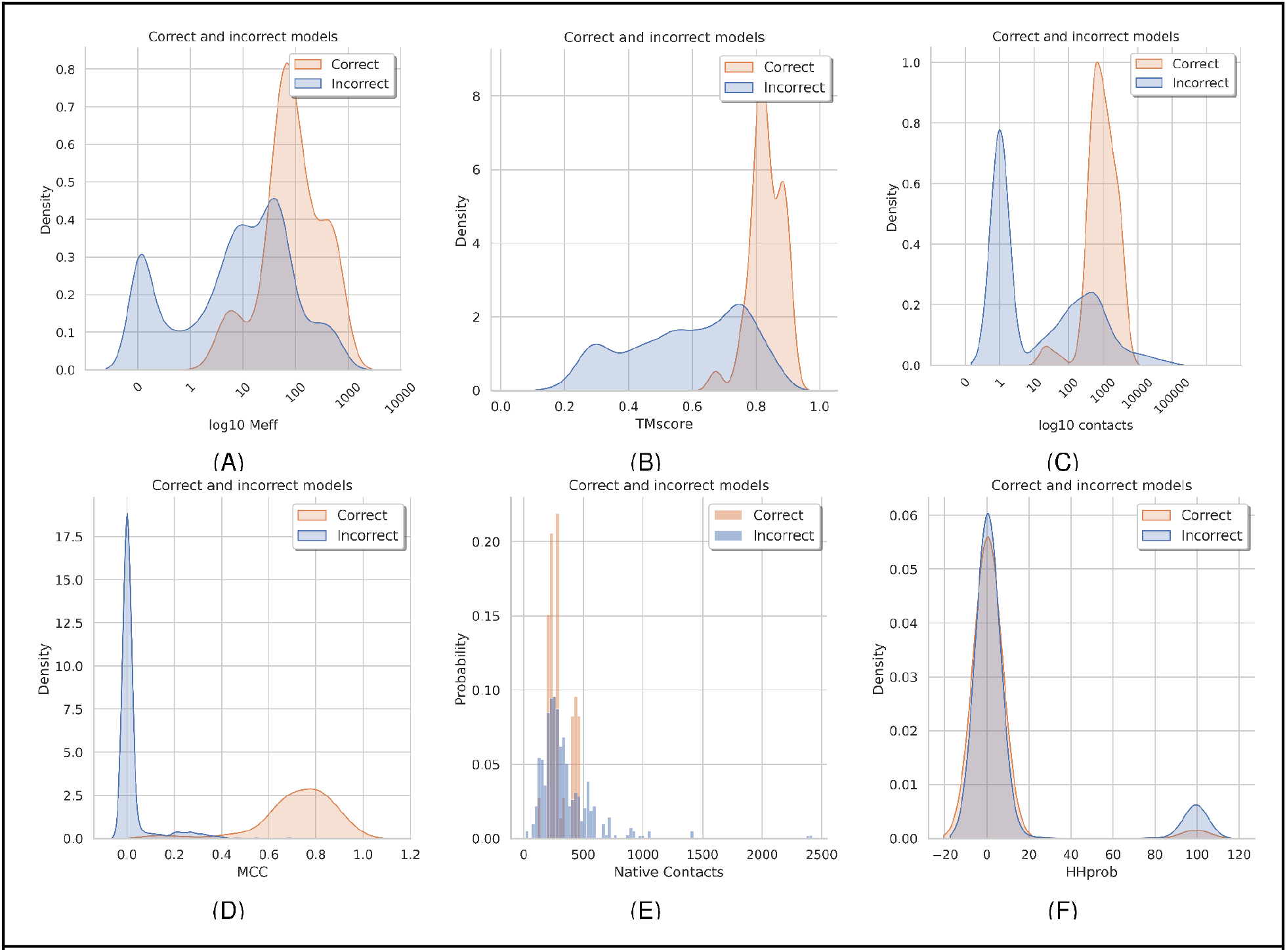
Features separating good and bad models. Good models are defined as having a dockQ score over 0.23. All models from all the methods marked with a star in table 1 are included. A) Distribution of the number of effective sequences (log10 of Meff), (B) average TM-scores of chain A and B, (C) number of inter-chain contacts predicted, (D) MCC values describing the accuracy of the inter-chain contacts, (E) number of contacts in the native interface, and (F) HHprob (similarity) between chain a and chain b in the complex.

First, it can be seen that the successful dockings tend to have a multiple sequence alignment of one hundred or more residues, see Figure 5A. It can be noted that about 25% of the protein pairs have less than ten effective sequences in the merged alignments and 50% less than 100. Only three proteins with 100 sequences or less have a correct model and no one with less than 35 effective sequences. The TM-score of all correctly docked models is high (>0.67), not surprisingly, as TMscore increase with larger MSAs. However, there are also many models with good TM scores but with incorrect docking. Interestingly, many good single chain models with very few sequences in the MSA exist. The highest TM-score for a model with only one sequence in the MSA is 0.56.

Next, when studying the number of inter-chain contacts predicted, it is clear that there is a narrow range of contacts around 100 (average 125) in all successful models. These predictions are mostly correct (MCC values over 0.5). In contrast, a large set (50%) of all models have no contacts predicted. However, many incorrect models have a similar number of predicted contacts as the correct models, but these predictions are simply wrong (MCC values close to zero). Interestingly, some unsuccessful models have more contacts predicted than the successful ones, and most models (both correct and incorrect) have about 500 contacts (<12Å) in the native structure.

### Pseudo homodimers and repeats can cause artefacts

As noted above, some unsuccessful models have very many inter-chain contacts predicted, caused by artefacts similar to the one shown in Figure 6A-D. This type of artefacts seems to be caused by homology between the two chains (sequence identity of 29% for 3qlu). This homology generates the coevolutionary signal from the intra-chain contacts to be reproduced as inter-chain contacts. The protein 3pv6 (sequence identity 29%) is even more complicated as the artefact does not cover the entire first chain. Still, instead, the first chain contains two homologous domains and the second chain a third homologous domain. Another type of artefact is seen in 4yoc, where a larger number of incorrect predictions occur.

**Figure 6.**
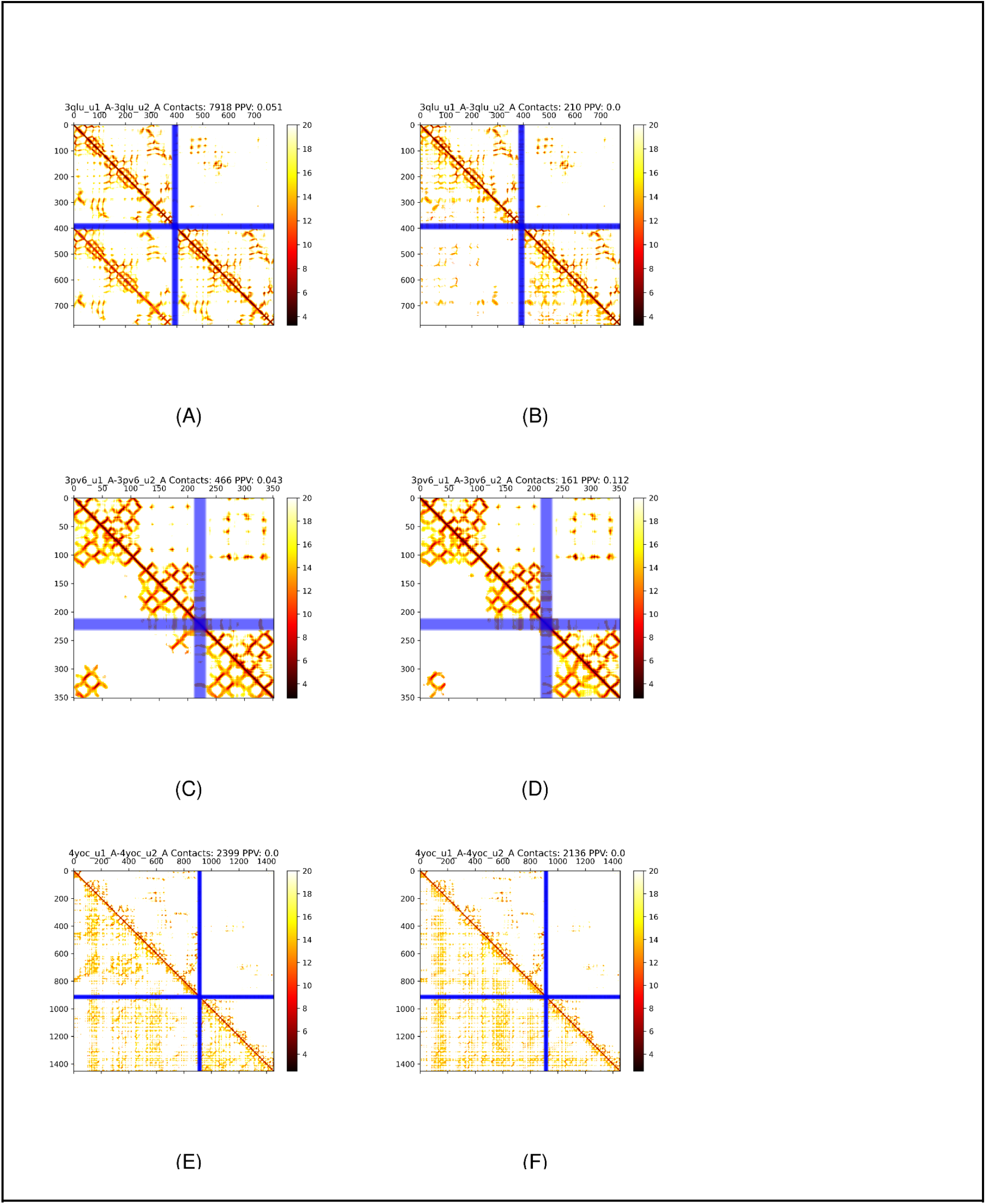
Distance maps for predicted (lower left) and native (upper right) distances for three proteins with artefacts 3qlu (A,B), 3pv6 (C,D) and 4yoc(E,F). Left columns (A,C,E) with N3 alignments, right (B,D,F) with reciprocal best hits.

To study how similarity between the two chains affect the docking results, we used HHalign to compare the multiple sequence alignments of the two chains in a complex. In Figure 5F, it can be seen that a set of (21) protein pairs exist with an HHalign probability of 99% or higher. The vast majority of these models are wrong (19), and some (9) of them have more than 1000 predicted contacts. We tried different strategies to reduce these artefacts, including reciprocal best hits, Figure 6. However, when this is done, the actual inter-chain contacts are also lost. It is possible that other strategies, similar to those used for homodimers [37], could work better. Still, we have not succeeded in getting rid of the artefacts without losing the actual contacts.

### Consensus scoring can separate correct and incorrect models

Is there a possibility to distinguish between correct and incorrect models? When modelling individual proteins, it has often been useful to compare the produced models to measure their reliability. These types of quality estimates have been successful in CASP since CASP5[40,41]. Here, we estimate to use the same idea for docking.

For consensus scoring, it is necessary to have at least two models to compare and measure the similarity of models. We examined two alternative sets of models and two alternative scoring functions. We can compare the two alternative orientations of the merged alignments (chain A-B vs B-A) or generate five models with the same orientation. The models can then be compared using dockQ or MMscore[38], see Table 2.

**Table 2:** Overview of consensus methods used to rank models.

Consensus scoring can separate correct and incorrect models.
| Name | Comparison method | Models to compare |
| --- | --- | --- |
| PconsDock-MMpair | MMscore[38] | Reversed alignments |
| PconsDock-MMcons | MMscore | Five models |
| PconsDock-dockQpair | dockQ[39] | Reversed alignments |
| PconsDock-dockQcons | dockQ | Five models |
| Table 2: Overview of consensus methods used to rank models. |  |  |

All four methods are excellent at separating the correct and incorrect models (AUC>0.93) for all methods. However, the methods that use five models are slightly better, Figure 7, possibly due to the (few) cases where only one of the contact maps generates a good model, as in the reverse order maps, both models have the same score. All four consensus methods identify a few more correct methods than the models from the best single alignment, and the MMcons and dockQcons methods also identify one or two more correct models compared to dockQpair and MMpair, Figure 7A.

**Figure 7:**
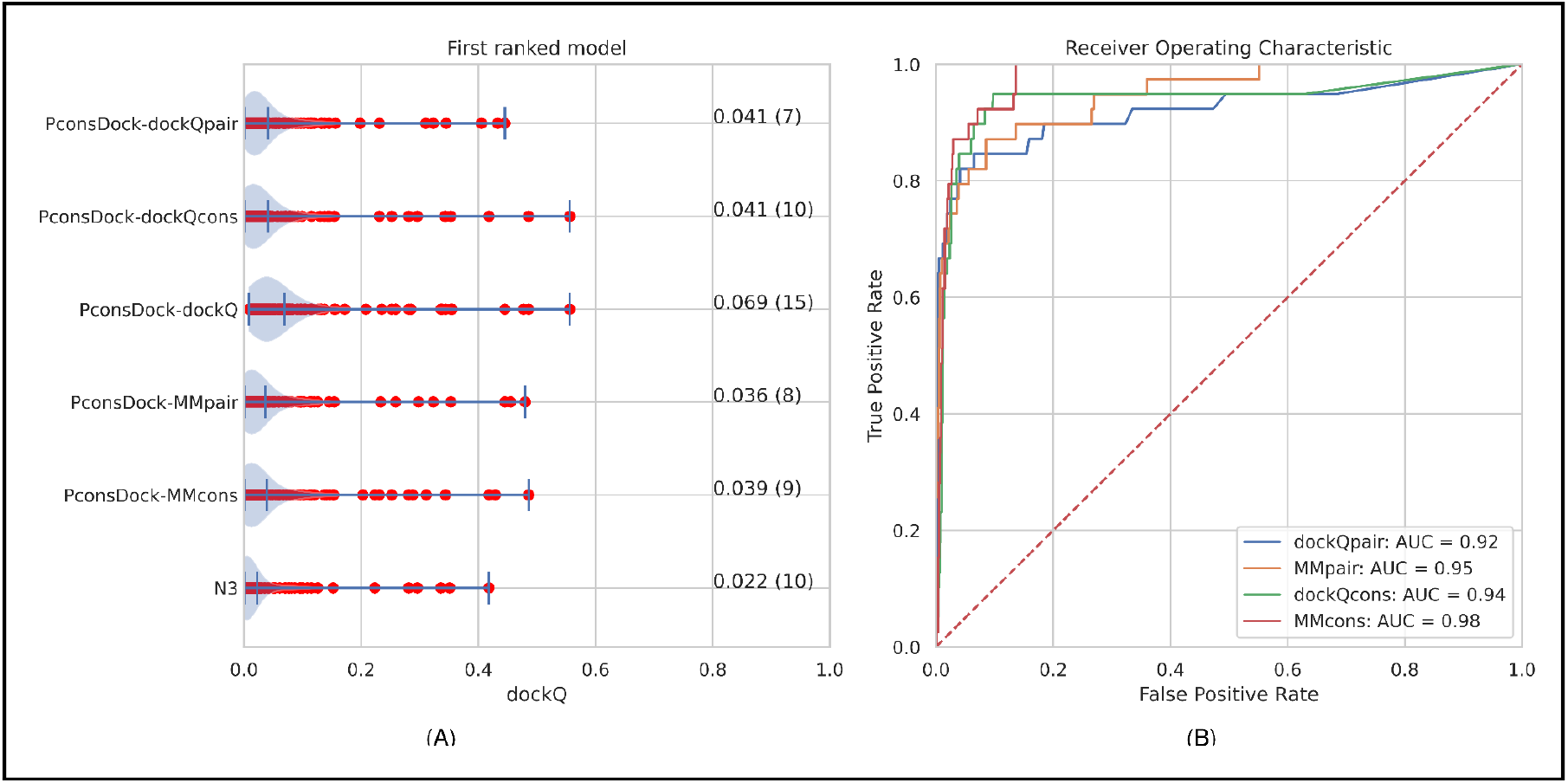
Different methods to identify the best possible model from models generated with different alignments. A) dockQ scores for first ranked models selected by a consensus approach using four different methods (see Table 2) compared with best possible choice (PconsDock-dockQ) and using a single method (N3). B) Receiver operator curve for separating correct and incorrect models using the four different consensus methods.

### Successful models for all kingdoms of life

The basic information used to dock two proteins here is coevolution in conserved interaction patterns in protein-protein interfaces. For detection of these signals, it is necessary to identify the protein pairs that interact in the same way, and this is much easier if there are few (or no) paralogs and the proteins only are involved in a few (or only one) specific interaction. Presumably, this is easier for prokaryotic protein pairs as these have fewer paralogs and, therefore, the interaction partners are more likely to be conserved among the identified orthologs. However, it is also possible that using both eukaryotic and prokaryotic sequences can help[42].

Out of the 15 models with successful predictions, the majority are predominantly unique to bacteria (as defined with more than 75% of the sequences in the merged alignment being bacterial), Figure 8A. However, four are mixed, as defined that no kingdom has more than 74% of the sequences, three are mainly (>75%) eukaryotic, and one consist mainly of archaea. The fact that successful prediction exists in all classes shows that this methodology is not exclusively useful for bacterial protein pairs, although the performance is, on average, slightly better for the prokaryotic protein pairs, Figure 8A.

**Figure 8.**
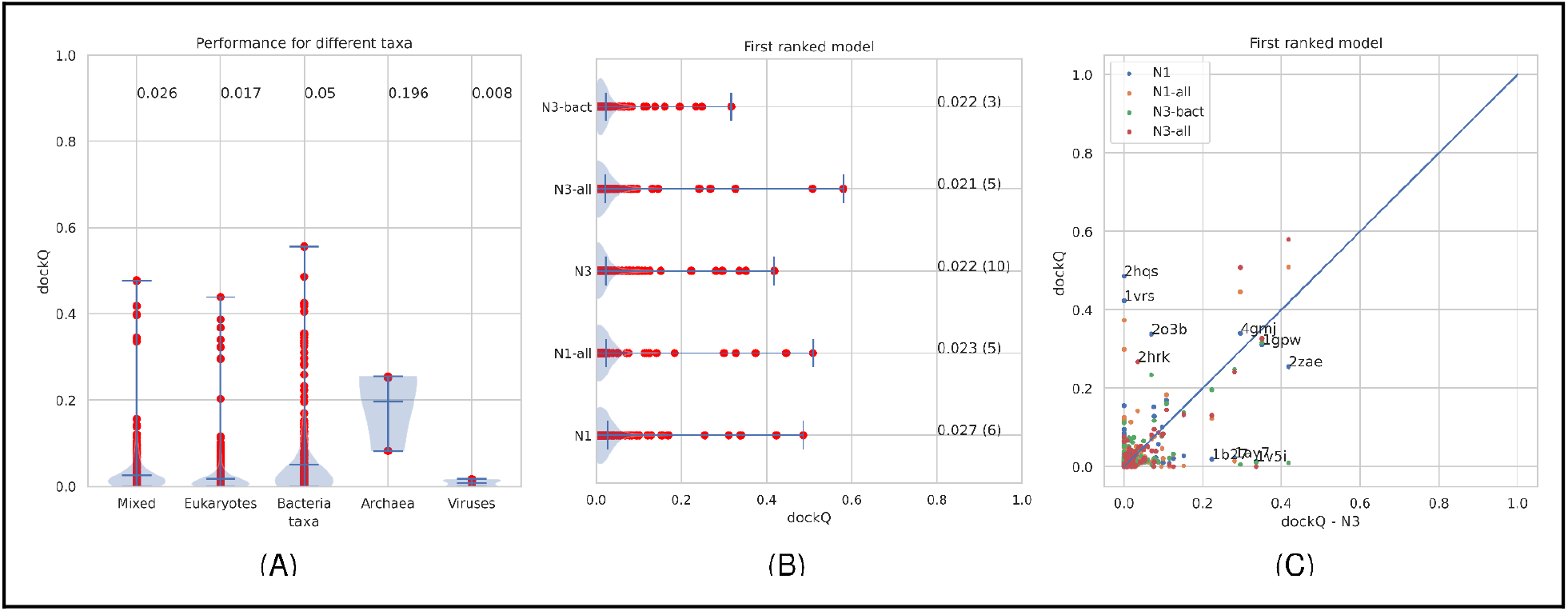
A) Prediction qualities for all models (using the methods to produce an alignment marked with a star in Table 1). B) Predictions qualities for all models using different alignments and sequence databases (see Table 1 for details). C) Comparison of dockQ scores for individual models compared to the N3 models.

Next, we tried to use a bacterial specific sequence database for constructing the multiple sequence alignments (N3-bact), Figure 8B and C. Here, it can be seen that the quality of a few models got worse, and only one improved significantly. The dockQ score of 2o3b (Nuclease A from E.coli) improved from 0.00 to 0.23. The paired multiple sequence alignment for 2o3b when using all reference proteomics is mainly eukaryotic, showing that in this case, the inclusion of eukaryotic genomes generates noise, making the signal getting lost. It also shows that a smaller, Meff=68 vs 306) multiple sequence alignment is sometimes to prefer. One example where the signal is lost when eukaryotic proteomes are excluded from the MSA is 2zae (archaeal homolog of the human protein complex Rpp21-Rpp29), whose dockQ scores drop from 0.42 to 0.01. In the original alignment, about 70% of the sequences are eukaryotic, and the size of the MSA drops from 839 effective sequences to 1 effective sequence.

In the alignments discussed above, we have only used the reference proteomes, but it is also possible to use all complete proteomes from UniProt. This dataset is more than three times larger, and most of the additional proteomes are bacterial. Figure 8 shows that the overall performance does not change significantly by using the larger database. However, there are a few targets whose performance increases significantly. The most striking improvement is for 2zae, whose dockQ score increases from 0.43 to an impressive 0.58 (Fnat 0.468 iRMS 1.975 LRMS 2.745 Fnonnat 0.326). The prediction of 2zae is the best prediction obtained. Another example is 2hrk (Arc1p and MetRS from yeast) which improves from 0.07 to 0.32. Anyhow, the inclusion of many more proteomes only makes a significant impact on a few proteins.

We also examined the subcellular location of the successful targets. There exists a certain tendency to have more membrane related (4/15 compared to 9% in the dataset) interactions. The successful targets include periplasmic (3/15), extracellular (3/15), cytoplasmic (4/15), and one nuclear target. Given the low number of successful docking cases, we can not judge the significance of any preference for any specific localisation; anyhow, the methodology can be applied to targets from various locations.

### Comparison to TMdock and GRAMM

How well does the fold-and-dock methodology compare with traditional docking methods? First, we compared it to one shape complementarity method, Gramm, and one template-based docking method, TMdock (see Figure 9. In the pure numbers, it can be seen that the FFT-based GRAMM with the AACA18 potential for ranking outperforms the other methods. Further, using the contacts (either from trRosetta or RaptorX) as a scoring potential does not improve the performance of GRAMM, rather than the reverse. Because of the exclusion of templates similar to targets (see Methods), TMdock performs worse than the other methods.

**Figure 9:**
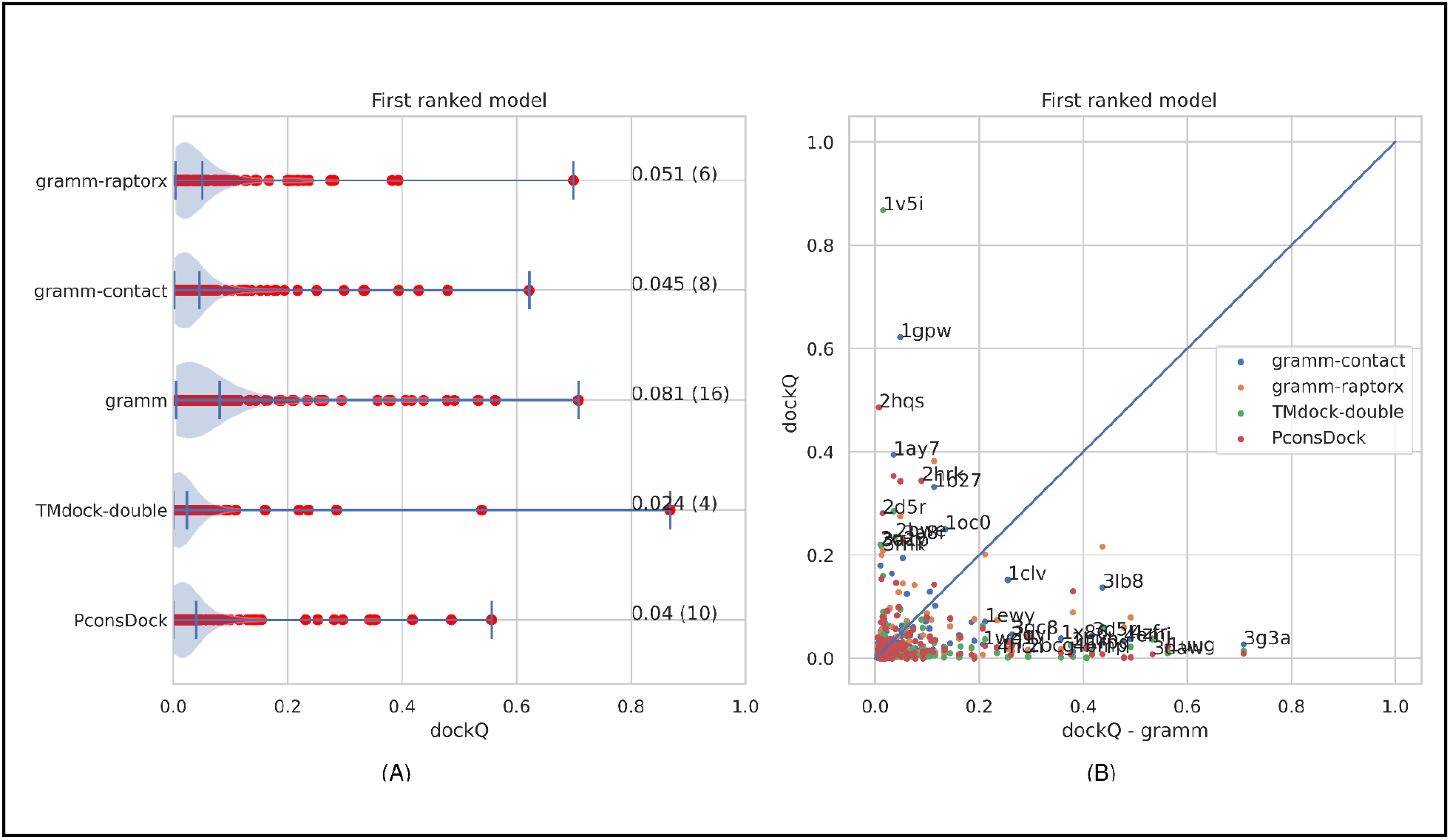
Comparison with GRAMM and TMdock. Gramm-contact is the gramm-scores using predicted contacts as a constraint, gramm is gramm-docking ranked by the AACE18 scoring potential, TMdock-double is a template-based docking method.

In summary, none of the docking methods can be seen as wildly successful as all methods predict less than 10% of the first ranked models correctly. However, there is room for improvement as the results appear complementary, Figure 9B. No single model that GRAMM accurately predicts is accurately predicted by PconsDock or vice-versa. Therefore a combined method could, in the future, be used to improve the performance.

### CASP14 -successful prediction of H1065

We used the PconsDock approach to predict intra-and inter-distance contacts using trRosetta for all relevant targets in CASP14. For a few models, we got exciting fold and dock results; see the contact map in Figure 10. These contacts were then used as a guide for further analysis and additional tests. In most models, this approach did not perform better than other methods, but for one model (H1065), our third-ranked model was ranked as the best of all models submitted to CASP14. The third model was generated by TMdock [43], refined using Rosetta minimisation. Input monomers were selected from the models produced after the CASP modelling stage 2, considering ProQ4 scores and visual inspection. Submitted docking was obtained by running the best server models, selected by ProQ4, using TMdock against a library of interface-only structures. The selected model was the model most resembling the first ranked model from the fold-and-dock approach. According to the official CASP evaluation, the MMscore of the three models are 0.60, 0.48 and 0.84, and the global QS scores 0.092, 0.060 and 0.685, clearly showing that the third model is better than the others. However, the structural similarity between the third and the first ranked model (generated directly by PconsDock) is high. This success shows a potential path for further improving the results of a fold-and-dock is to include an additional refinement protocol.

**Figure 10:**
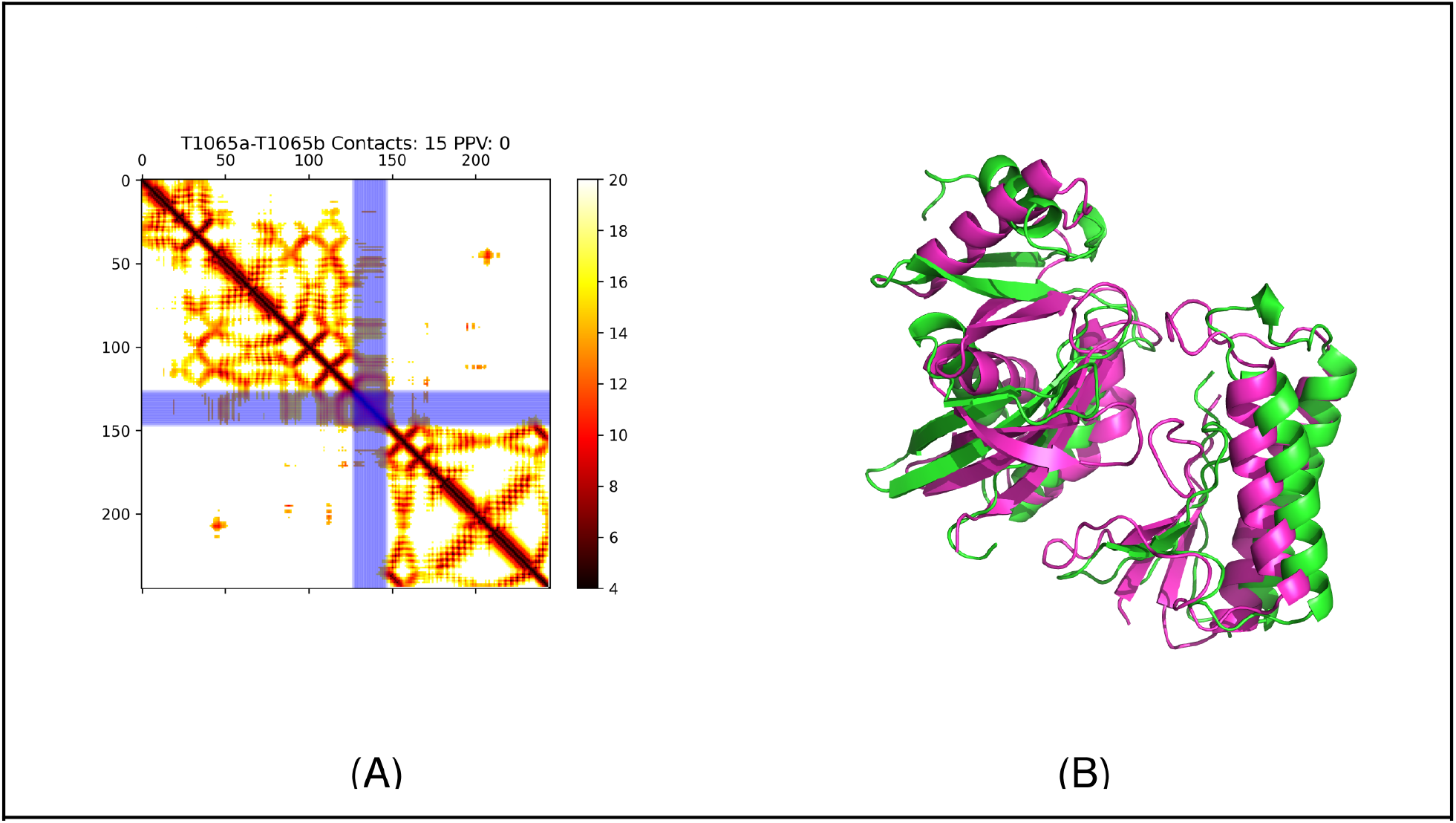
A) Predicted contact map of CASP target H1065, B) Models of H10165, green model (fold and dock) and magenta model generated by TMdock using the PconsDock model for selection). The structure of H1065 is not available for the public yet, so we can not perform a direct comparison with the native structure.

## Conclusions

Here, we present an analysis of a fold and dock protocol, PconsDock, based on predicted intra-and inter-chain distances using trRosetta. We show that it is possible to produce acceptable models using the fold-and-dock protocol for some targets. The success rate is comparable to traditional docking methods. However, we do believe that the potential of this type of fold-and-dock protocol is more extensive, as the method is not dependent on the structure of the individual protein chains., i.e. fold-and-dock methods are applicable to predict the structure of all complexes, including the ones involving flexible proteins.

One limit of this study is that the network used here (trRosetta) is trained to predict intra-chain contacts, but we use it for predicting inter-chain contacts. We tried to train a network specifically to predict inter-chain contacts, but the performance was not as good as trRosetta, possibly because it was trained on a significantly lower number of contacts. Neither is the RaptorX methodology specifically trained to predict inter-chain contacts significantly better than trRosetta.

The problem that remains unsolved is the choice of the best alignment for folding and docking. In some cases, only specific alignment gives correct folding and docking based on the intrinsic evolutionary characteristic of the proteins and their interaction. Therefore, we believe that improving this methodology lies in improving merging the two multiple sequence alignments. Identifying *a priori* the alignment containing more information is still an open challenge.

## Material and Methods

### Dataset

To evaluate the performance of various docking methods, we use the unbound structures for 221 hetero-dimeric only protein complexes from Dockground 4.3 [18], as we can not use the fold-and-dock protocol for homodimers (see the figshare repository [44]).

### Evaluation

The main evaluation criteria to evaluate the success in docking used here were the dockQ score [18,39], which gives 0 to a random prediction and 1 to a perfect prediction. Here, it should be noted that a dockQ score over 0.23 roughly corresponds to an “acceptable” model in CAPRI [45], and we will therefore call all models with dockQ >0.23 as correct and all others as incorrect. To evaluate the quality of the individual models, we have used TM-score [45,46]. We also MM-align for comparing docked models [38].

We do also analyse the accuracies of the distances predicted by trRosetta with actual distances. For simplicity, we have in several cases redefined the distances (both predicted and real) as contacts by using a cutoff of 12 Å, i.e. only predicted or native distances shorter than 12 Å are included. Here, the probability distance distribution from trRosetta is converted by using the weighted means of all probabilities.

### Paired alignments

The critical component in our algorithm is the formation of paired alignments, i.e. a set of aligned protein pairs assumed to interact in the same way. Starting from two proteins, which are assumed to interact, we search both sequences against a proteomic database using jackhmmer [27,38]. Several different alignment parameters and databases were tried; see Table 1. The default database used is all reference proteomes from UniProt [47] as of May 2020. This dataset consists of 55 million sequences. In addition, we tried to use only bacterial proteomes (30 million sequences) or all (non-excluded) proteomes from UniProt (199 million sequences) with a few alignment parameters, see above.

The next step is to create a paired alignment for the two protein chains. Here, for each protein where both chains had a significant hit (over the defined threshold), the top hit was included unless both proteins had identical top hits. In such cases, that proteome was ignored. We also tried to use a reciprocal best hit, i.e. only including the top hits if the original proteins also were top-ranked for these proteins in their proteome.

When paired sequences are identified, they are merged to form a paired multiple sequence alignment. Here, 20 glycines are inserted between the two sequences to avoid edge effects. The two alignments can be merged in two different orders, and both were tried as in a few cases, one of the orders provided better predictions.

Finally, the paired alignment was “trimmed” to take away sequences with too many gaps. By default, sequences with more than 25% gaps in the merged alignment were excluded, but other parameters were also tried (see Table 1).

### Distance predictions

Distance and angles were predicted using trRosetta from the paired alignment. We also tried one alternative method to predict contacts, RaptorX using the protocol for predicting complex interactions [36]. However, this method does not provide distances, just contacts, and therefore, it is necessary to add predicted secondary structures from psipred when using these contacts. The distances were then used in Rosetta as described in the original trRosetta protocol.

### The fold and dock protocol

Fold and dock were performed using the same protocol and constraints as in trRosetta for the two chains separated (obviously, they were treated as two separate molecular objects). The same optimisation protocol (minmover) was used. However, an additional set of inter-chain constraints were added in the form of a weak flat-harmonic potential between all inter-chain pairs of residues with a predicted contact probability of over 50%. These constraints were necessary to ensure that the two chains were modelled near each other; for details, see the GitHub repository. We also used pyconsFold [35] for fold-and-dock, both with distances predicted by trRosetta and with contacts predicted by RaptorX.

### Shape complementarity docking

For comparison, we used the GRAMM scan stage [12,35] with the default parameters in order to generate an initial set of docking decoys for the same dataset. For ranking the decoys, we used the AACE18 contact potential [16]. In addition, we used the predicted contacts (all with probability > 0.5 and distance shorter than the predicted distance plus 2Å) as a constraint to GRAMM. We used both trRosetta and RaptorX to predict the distances.

### Template-based docking

We also used the TMdock [13] with the standard full-structure template library for template-based docking. To avoid including templates with high sequence identity to the target structure, we excluded all template hits where both chains in a complex have significant(E-value < 10^−2^) similarity to the templates. If this is not done, the performance of TMdock would be much higher.

## Availability

All scripts for predictions and analysis are available from

https://github.com/ElofssonLab/bioinfo-toolbox/trRosetta/

Details for each run is available from

https://github.com/ElofssonLab/bioinfo-toolbox/benchmark5/benchmark4.3/.

All models joined alignments, and evaluation results are available from a figshare repository[44].

